# BERT-T6: Towards High-accuracy T6SS Bacterial Toxin Identification Using Protein Language Model

**DOI:** 10.1101/2025.10.17.683028

**Authors:** Xianwei Mo, Jianxiu Cai, Shirley W. I. Siu

## Abstract

Type VI secretion system effectors target the cell wall, membranes and nucleic acids, leading to the killing of bacteria or impairment of host cell defense mechanisms. Accurate identification of T6SEs will be beneficial to understand the virulence of these bacteria via type VI secretion systems as well as bacterial pathogenesis. Although some traditional machine learning-based and deep learning-based tools have been developed to distinguish T6SEs from non-T6SEs, we believe there is still room for further improvement. To obtain the robust feature for model construction, we successively investigate various classic sequence-based features and embeddings from pre-trained transformer-based protein language models. Building upon the model incorporating ProtBert embeddings, we employed a transfer learning approach to fine-tune the ProtBert protein language model with a downstream T6SE classification task. The resulting BERT-T6 model demonstrates performance significantly superior to baseline models. More importantly, with an accuracy of 0.959, a sensitivity of 0.909, a specificity of 0.973, a precision of 0.905, a F1-score of 0.907, MCC of 0.881, our model achieves performance competitive with state-of-the-art binary and multi-class predictors. This work highlights the effectiveness of utilizing BERT with transfer learning for T6SE prediction. BERT-T6 provides a robust and precise approach for identifying T6SEs, offering promise for enhancing studies of bacterial virulence mechanisms.

## 1. Introduction

Type VI secretion systems (T6SSs) are a versatile nanomachine found to distribute in more than 25% of the Gram-negative bacteria including most pathogenic ESKAPE strains, namely, *Enterococcus faecium, Staphylococcus aureus, Klebsiella pneumoniae, Acinetobacter baumannii, Pseudomonas aeruginosa and Enterobacter* species, which are responsible for approximately 75% of infections and deaths by antibiotic-resistant bacteria [1–4]. By means of T6SS, the effector proteins are injected into eukaryotic or competitor bacterial cells, playing an important role in both bacterial pathogenicity and microbial competition[5]. Therefore, identifying T6SS effectors (T6SEs) is helpful for understanding the virulence of bacteria via T6SS and developing new therapeutic strategies.

Experimental methods for the discovery of T6SEs rely mainly on specific analysis of T6SS-associated genes, proteomics-based methods and screens of mutant libraries, which are costly and have low identification rates due to the versatility and uncertainty of protein expression and secretion levels. Meanwhile, some conserved domains have been found to be closely related to T6SE and can be utilized to identify and discover new T6SEs, including Rhs/YD repeat sequences, proline-alanine-alanine-arginine (PAAR), TTR domains and MIX motifs[6–8]. However, these bioinformatics methods heavily depend on existing knowledge of T6SEs, their sequential features and biochemical properties, which make it a challenge to recognize T6SEs having novel sequence pattern.

Machine-learning method, including deep learning, are considered as promising strategies to discover novel T6SEs. Sequence feature-based machine learning models were first developed to accurately predict T6SS effector proteins. These models are constructed based on manually selected features, and tend to complete the prediction through voting strategies, which might be beneficial to address poor performance caused by the imbalance of dataset[9, 10]. However, most sequence-based features, which carry limited information, generally cannot effectively distinguish T6SEs from non-T6SEs. As effectors with different sequence patterns being continuously discovered, previous manually selected features may be no more suitable for model construction. In addition, the redundancy of sequence adversely affects model generalization.

The emergence of protein language model (PLM) has made the automatic extraction of protein features possible. Without any experimental annotation, PLMs are typically trained on hundreds of millions to billions of sequences using self-supervised neural networks. The evolutionary and sequential information can be effectively extracted and ultimately described as embedding from the last hidden layers, which has been widely used for downstream protein-related prediction tasks. Given the limited scale of the dataset of T6SEs, universal features generated from PLMs cannot represent T6SEs well. Therefore, some studies tend to adopt an approach that combines sequence-based features with protein language model-derived features to construct predictive models[11]. However, these models still critically depend on manually engineered features and lack the flexibility to dynamically capture the semantic information of T6SEs as the dataset scale expands, ultimately leading to less effective prediction performance[12]. Transfer learning is a machine learning technique that adapts generalized protein features for a specific task by fine-tuning a pre-trained model on a smaller, task-specific dataset, thereby enhancing prediction performance. This approach has been applied in the prediction of signal peptides[13], protein secondary structure[14], protein stability[15], protein-protein interactions[16], protein-coding variants[17] and antigen specificity[18].

Inspired by the above, here, a step-by-step approach was adopted to explore the model used for T6SE prediction. And we successively investigated the classic sequence-based features, pre-trained PLM-based embedding and BERT-based transfer learning approach for model construction. Ultimately, we proposed the model BERT-T6, which, compared with the previous two types of models, improves performance and mitigates the imbalance between sensitivity and specificity by fine-tuning the ProtBert PLM on downstream T6SE classification tasks. Importantly, BERT-T6 achieves competitive performance with state-of-the-art (SOTA) predictors on test set, especially in the accuracy, sensitivity, F1-score and Matthews’s correlation coefficient. Given the predictive power and interpretability of the transformer model, BERT-T6 will facilitate the discovery of novel T6SEs and enable studies of their functional involvement in the pathogeneses of bacterial diseases.

## 2. Materials and Methods

### 2.1 Data collection and preparation

The dataset of experimentally verified T6SEs was constructed through manual collections of the corresponding databases, which includes 138 T6SEs from Bastion6[9], 329 T6SEs from SecReT6[19], and 111 T6SEs from TxSEdb (http://61.160.194.165:3080/TxSEdb/index.html). And the negative samples were used the same as Bastion6, containing 1112 non-effector sequences. Then CD-HIT[20] was employed to remove homologous-redundant sequences with sequence identify > 60%. Eventually, we obtained a non-redundant training dataset containing 305 positive and 1109 negative protein sequences.

To further evaluate the performance of our approach, we constructed an independent test set. The sequences in this set are sourced from the independent test sets from literature[9, 10] and newly experimentally validated T6SEs from recent papers[21–34]. After these sequences were preprocessed with CD-HIT at a 60% threshold to remove highly homologous samples, we obtained a final set of 59 T6SEs and 200 non-T6SEs.

### 2.2 Feature extraction

In this study, two types of features were extracted for construction of classification model. They are classic sequence-based features and embeddings from pre-trained transformer-based PLMs. About 23 sequence-based features were extracted using *iLearnPlus* Python package[35], and they are amino acid composition (AAC), composition of k-spaced amino acid pairs (CKSAAP), di-peptide composition (DPC), dipeptide deviation from expected mean (DDE), tri-peptide composition (TPC), grouped amino acid composition (GAAC), composition of k-spaced amino acid group pairs (CKSAAGP), grouped dipeptide composition (GDPC), grouped tripeptide composition (GTPC), Moran, Geary, normalized moreau-broto autocorrelation (NMBroto), CTDC, CTDT, CTDD, conjoint triad (CTriad), conjoint k-spaced Triad (KSCTriad), sequence-order-coupling number (SOCNumber), Quasi-sequence-order (QSOrder), pseudo-amino acid composition (PAAC), amphiphilic pseudo-amino acid composition (APAAC), adaptive skip dinucleotide composition (ASDC), PseAAC of distance-pair and reduced alphabet (DistancePair).

The pre-trained transformer-based model used for generating protein embedding consists of Evolutionary Scale Modeling (ESM) family including various ESM-2 with dimensions of 320, 480, 640, and 1280, ESM-1b and ESM-1v, BERT-based model including ProtBERT and ProtBert-BFD, trained on UniRef100 dataset or Big Fantastic Database (BFD), T5-based model including ProtT5-XL-UniRef50 and ProtT5-XL-BFD, which also trained on UniRef50 dataset or Big Fantastic Database (BFD), Albert-based model (ProtAlbert) trained on UniRef100 corpora. The publicly available code and pretrained ESM models were obtained from ESM GitHub repository (https://github.com/facebookresearch/esm), others were sourced from the ProtTrans GitHub repository (https://github.com/agemagician/ProtTrans). Then protein embeddings (features) were extracted from the last layer of PLM utilized for subsequent machine learning modeling. The protein embeddings represented as a 2-dimensional array with a size of words embedding size × length of sequence. Ultimately, each protein or peptide sequence can be encoded as features with the same length as words embedding size after mean pooling in the direction of sequence length.

### 2.3 Machine learning methods for T6SE prediction

To compare with BERT-based transfer learning approach, various machine learning algorithms were investigated for constructing T6SE classification models using the *Scikit-learn* Python package. These algorithms included Random Forest (RF), Support Vector Machine (SVM), Multi-layer Perceptron (MLP), K-Nearest Neighbors (KNN), Bagging, Decision Tree (DT), Logistic Regression (LR), AdaBoost, Gradient Boosting Decision Tree (GBDT), Linear Discriminant Analysis (LDA), and Naive Bayes. In addition, the XGBoost algorithm is implemented in the xgboost Python package, while LightGBM is implemented in the lightgbm package. Default values from *iLearnPlus*[35] were used for the hyperparameters of the machine learning algorithms above.

### 2.4 BERT-based transfer learning approach for T6SE prediction

The BERT architecture, whose full name is Bidirectional Encoder Representations from Transformers, is an encoder-only structure typically focuses on representing languages. This architecture consists of multiple stacked transformer blocks, where each encoder layer employs multi-head self-attention followed by feedforward neural networks, all trained using masked language modeling. This method masks a part of the input for the model to predict. Additionally, the input contains an additional token, the [CLS] or classification token, which is used as the representation for the entire input. The [CLS] token is conventionally employed as the input embedding when fine-tuning for downstream tasks like classification[36]. One major advantage of pre-trained BERT models is that most of the training work has already been completed. Fine-tuning them on specific tasks typically requires less computational cost and fewer labeled data. Additionally, BERT-like models generate embeddings at nearly every step of their architecture. From another perspective, BERT can also be viewed as a feature extractor.

Similar to natural language models, PLMs are obtained by training on extensive protein sequence corpora. Common PLMs primarily include autoregressive-based models, represented by ProGen and ProtGPT2, and autoencoder-based models, represented by the ESM series[37] (e.g., ESM-1b, ESM-2) and ProtBert/ProtT5. Here, we employed ProtBert[38], which is a BERT-based architecture pretrained on approximately 2.17 billion protein sequences from UniRef100. The model is from ProtTrans[39] and can be accessed through the Hugging Face Transformers API.

#### 2.4.1 Fine-tuning for the T6SE prediction task

Transfer learning was performed to fine-tune ProtBert and train the binary T6SE prediction model downstream while AdamW algorithm was used as the optimizer for model optimization. The model architecture is shown in the right side of **Scheme 1**. It should be noted that the [CLS] token, located at the sequence’s beginning, is designed not to bind to any specific word. During both pre-training and fine-tuning stages, it learns global information through the self-attention mechanism, serving solely as a global representation. Therefore, [CLS] token is extracted for supervised learning and subsequent prediction. Our prediction model for downstream tasks only contains one fully connected layer (FC1 with 512 dimensions) as hidden layer, which incorporates dropout regularization to prevent overfitting. And the output is further used for final T6SE prediction.

Our dataset has a positive-to-negative sample ratio of nearly 1:4, which could make the model biased toward the majority class. To address class imbalance in the dataset, focal loss was employed as the loss function, improving the model’s recognition of minority class samples by reducing the impact of easily classified samples[40]. Focal loss was developed based on the cross entropy (CE) loss for binary classification:

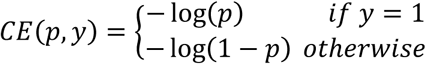

where y represents the ground-truth class and p is the model’s estimated probability for class 1, which is T6SE. For notational convenience, p_t_ is further defined as follow:

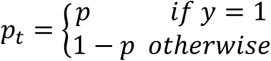

CE (p, y) is rewritten as CE (pt) = − log(p_t_). Eventually, the focal loss function is defined as follow:

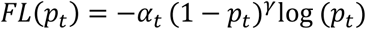

where α_t_ is a weighting factor used to balance class importance, (1 − p_t_)^γ^ represents a modulating factor, which reduces the contribution of easily classified samples to the loss and puts more focus on hard-to-classify samples[41].

**Scheme 1.**
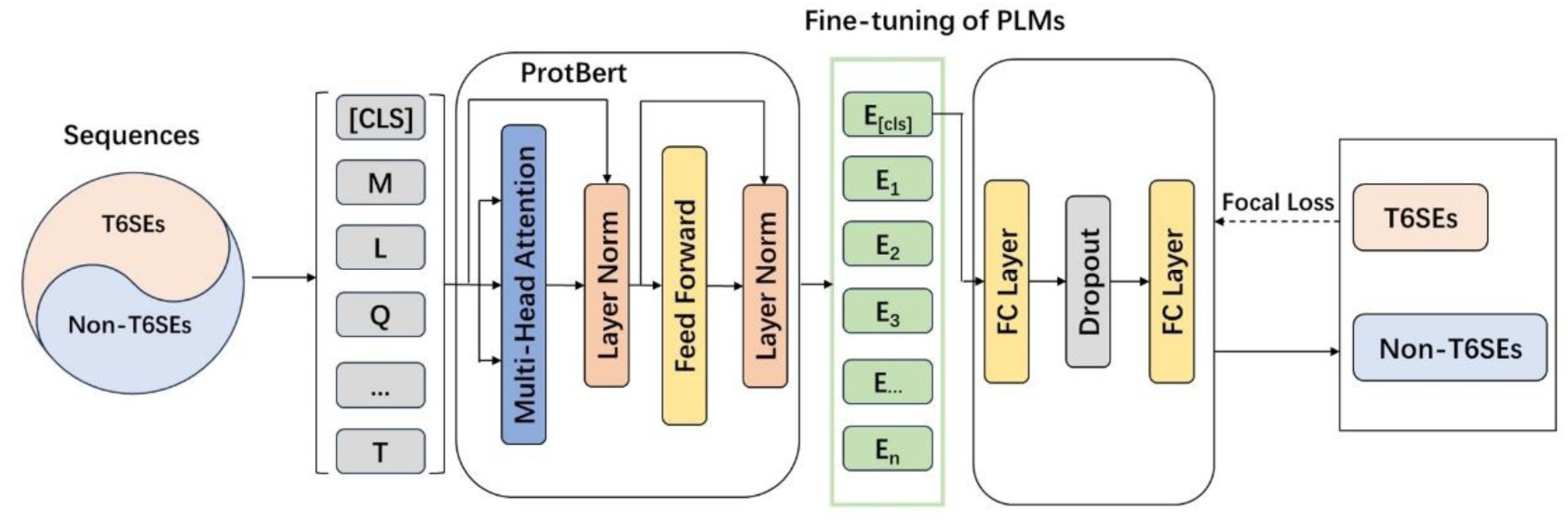
Illustration of the proposed BERT-T6 framework for T6SEs prediction.

#### 2.4.2 Hyperparameter tuning

To confirm the optimal parameters for model architecture, several hyperparameters were investigated. Initially, the performance of model with varying numbers of FC layers (1 to 4) were assessed under default parameter conditions. Subsequently, learning rate and dropout rate were investigated after determining the optimal number of layers. Lastly, various weight decay rates were tested for its influence on model performance. The candidate value range of each hyperparameters are exhibited in **Table 1**. During hyperparameter investigation, the data set was split in a ratio of 9:1 to generate training set and validation set. Therefore, the model was trained on 1272 sequences with a maximum epoch of 50 iterations, and validated on 142 sequences.

**Table 1.** Hyperparameters and candidate value ranges considered during model optimization.

| Hyperparameter | Candidate value | Optimal value |
| --- | --- | --- |
| number of FC layers | 1, 2, 3, 4 | 2 |
| weight decay rate | 0.1, 0.3, 0.5, 0.7, 0.01, 0.03, ...<br>1e-3, 3e-3, ..., 7e-3 | 5e-3 |
| dropout rate | 0.1, 0.2, 0.3, ..., 0.9 | 0.4 |
| learning rate | 1e-2, 1e-3, 1e-4, 1e-5, 1e-6 | 1e-5 |

After a series of experiments, the optimal hyperparameters were determined to be 2 FC layers, a learning rate of 1e−5, a dropout rate of 0.4, and a weight decay rate of 0.03, with a batch size of 6 after comprehensive consideration of several model performance metrics including sensitivity, F1-score, the area under the ROC curve and the area under the PRC curve. After all the hyperparameters were determined, the preprocessed dataset was randomly split into a training set (80%) and a test set (20%). Subsequently, the model was trained on the training set with the early stopping criteria and evaluated on the test set.

### 2.5 Model performance metrics

To quantify predictive accuracy and reliability, eight standard metrics were employed to comprehensively evaluate the performance of classification models, including Sensitivity (Sn), Specificity (Sp), Accuracy (ACC), Matthew’s correlation coefficient (MCC), Precision (PR), F1-score. These metrics offer complementary perspectives on classification performance, particularly in the presence of class imbalance. They are calculated according to the following formulas:

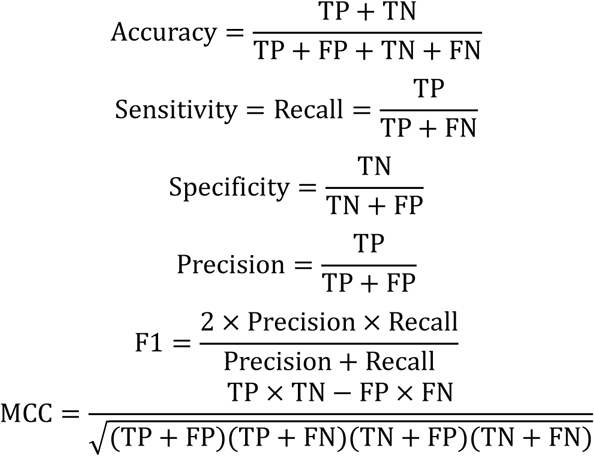

TN (True Negatives) refers to cases where negative samples are correctly identified while FN (False Negatives) represents positive samples that are mistakenly classified as negative. In addition to these discrete metrics, the models’ overall discriminatory performance was further examined through area under the receiver operating characteristic (AUROC) curves and precision-recall curve (AUPRC) values. This approach evaluates classification efficacy across the complete spectrum of decision thresholds, with higher AUC scores indicating more robust classification capabilities and superior generalization potential. Particularly for classification tasks with imbalanced datasets, the AUPRC is a more suitable and reliable evaluation metric than the AUROC.

### 2.6 Statistical analysis

Microsoft Excel and GraphPad Prism 8 were used for statistical analysis. The results are expressed as mean ± standard deviation (SD), while statistical significance was determined by the two-tailed Student’s t-test (ns, not significant; *P < 0.05; **P < 0.01; ***P < 0.001; ****P < 0.0001).

## 3. Results and Discussion

### 3.1 Peptide sequence analysis

To better understand the profile of data, length distribution and amino acid composition of T6SEs and non-T6SEs sequences were analyzed. As shown in **Figure 1**, the lengths of both T6SEs and non-T6SEs sequences have a wide distribution, covering a range from 0 to >1600. The majority of T6SEs sequences are 100-200 amino acids in length, while most non-T6SEs sequences range between 200-300. In addition, there is no significant difference in the amino acid composition between T6SEs and non-T6SEs, indicating that it is difficult to distinguish T6SEs from non-T6SEs by analyzing the frequency of key amino acids, which could otherwise reveal important information about the physicochemical properties of T6SEs.

**Figure 1.**
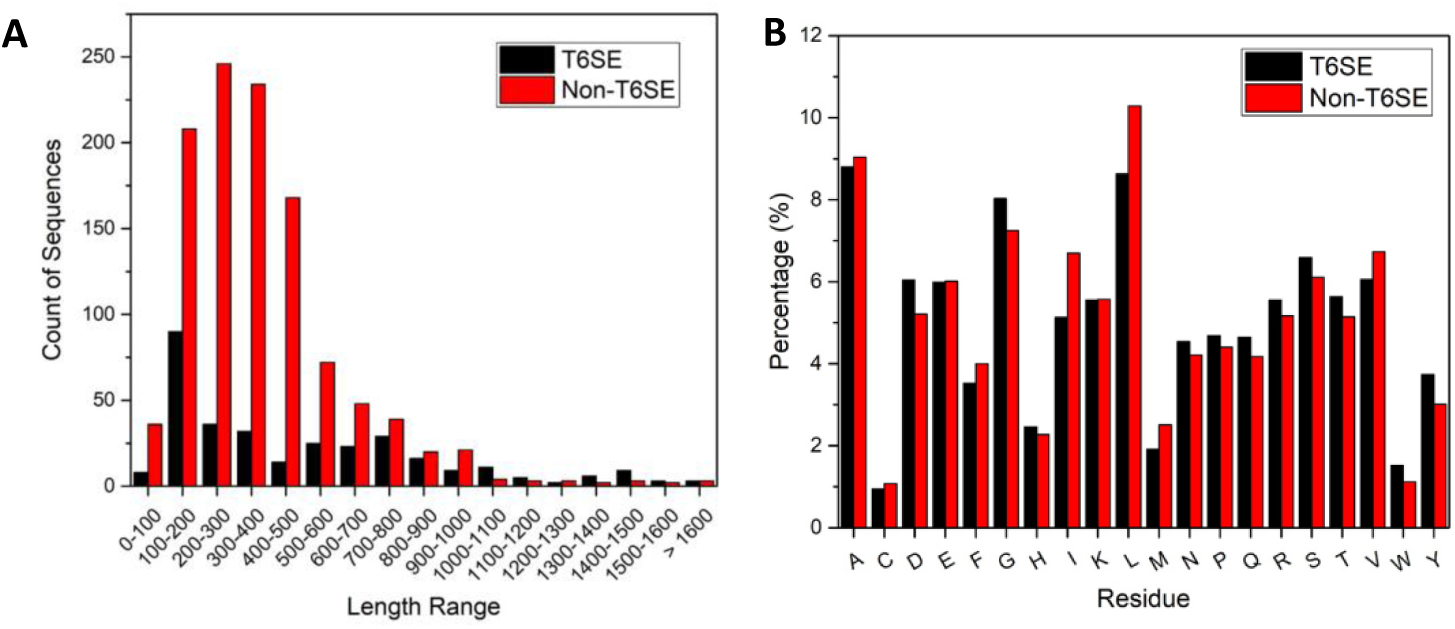
(A) Distribution of peptide sequence lengths for T6SEs and non-T6SEs. The plot illustrates the count of sequences across different length ranges. (B) Distribution of amino acid residues in T6SE and non-T6SE sequences represented as percentages. The bars show the relative abundance of each residue type among T6SEs and non-T6SEs, calculated from resides of 305 T6SE sequences and resides of 1109 non-T6SE sequences, respectively.

### 3.2 Models based on sequence-based features

The classic sequence-based features were first investigated for the construction of T6SE recognition model using various machine learning methods. Then splitting of training set and test set was the same as that described in “*2.4.2 Hyperparameter tuning*”. In addition, the robustness of model was evaluated using 5-fold cross-validation and predictive ability was evaluated on test set. To obtain the ideal feature for modeling of T6SE classification, SVM was employed as the default method. The performance of cross-validation is exhibited in **Table S1**, majority of models based on sequence-based features have a poor sensitivity with value of zero, indicating that they fail to recognize T6SE. Only models based on these five features have a sensitivity greater than 10%: DDE, QSOrder, CTDC, KSCTriad, and CTriad. Moreover, similar results can be observed in the evaluation on test set (**Table S2**). In total, model based on DDE features present the best performance whether it’s cross-validation or test set evaluation in terms of sensitivity, accuracy, F1-score, MCC, AUROC and AUPRC.

Next, DDE descriptors were set as default features to investigate various machine learning algorithms. Surprisingly, model based on SVM algorithm perform the best on both cross-validation and test set in terms of accuracy, MCC, AUROC and AUPRC (**Table S3** and **S4**). However, severe imbalance between sensitivity (66.13%) and specificity (95.05%) can be observed on results of test set evaluation, revealing the model cannot identify the positive samples well. That is mainly caused by imbalance of positive samples and negative samples, which requires further improvement. Eventually, the DDE_SVM model will serve as the baseline model for assessing the significance of BERT-based transfer learning approach on prediction improvement.

### 3.3 Models based on embeddings from pre-trained transformer models

The optimal model for predicting T6SE is obtained by testing combinations of pre-trained transformer models and downstream machine learning algorithms. The chosen PLMs for embedding and machine learning algorithms for generating predictions are shown in **Table 2**. The best machine learning algorithm to construct models based on different embeddings is not always the same (**Table 3**). For instance, the best machine learning algorithm for ESM-1b embedding based model is LR while for ProtAlBert embedding based model is MLP. In addition, it’s worth noting that MLP is an optimal algorithm for four types of embedding-based model, followed by SVM. In total, for the model based on ESM-2 embedding, with embedding dimension increases, the performance of model on test set has also gradually improved. However, the higher embedding dimension is not always means the better model performance for different PLM. The ProtAlBert model with the highest embedding dimension of 4096 performs not as well as ProtBert model with embedding dimension of 1024 on test set when employ the same MLP algorithm. Finally, the best model is a combination of features from the ProtBert model and MLP classifier, and achieves the highest performance on the test set. This model, named ProtBert_MLP, attains superior performance metrics, including Sn (0.919), ACC (0.965), MCC (0.897), F1 (0.919) and AUROC (0.948). The ProtAlBert_MLP model ranks second in terms of Sn (0.903), ACC (0.961), MCC (0.886), AUROC (0.940) and AUPRC (0.965). Additionally, ProtBert_MLP model also has the highest cross validation scores for Sn (0.881), MCC (0.853), F1 (0.884) and AUROC (0.976). Although ProtT5-XL-UniRef50_SVM model achieves the best results in another four metrics, i.e. Sp (0.986), PR (0.943), ACC (0.951) and AUPRC (0.943), it performs not as well as ProtBert_MLP model on test set. What’ worse, the imbalance between Sn and Sp can be observed not only during cross validation but also test set evaluation, and more significant than that of ProtAlBert_MLP and ProtBert_MLP model. Additionally, models based on PLM embeddings generally achieve better performance than those based on sequence-based features. Eventually, ProtBert was selected as the pre-trained model for fine tune using BERT-based transfer learning approach.

**Table 2.**
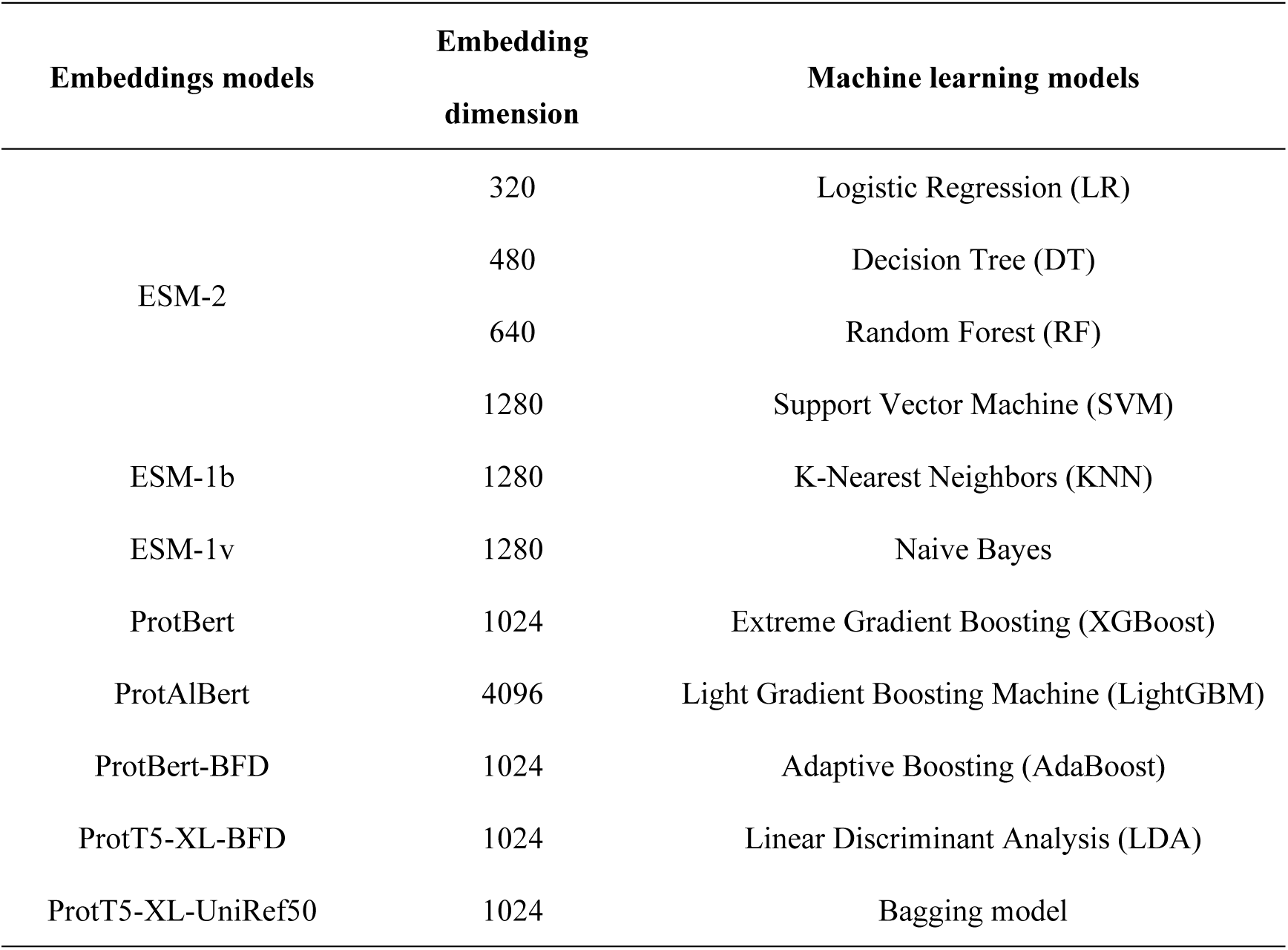
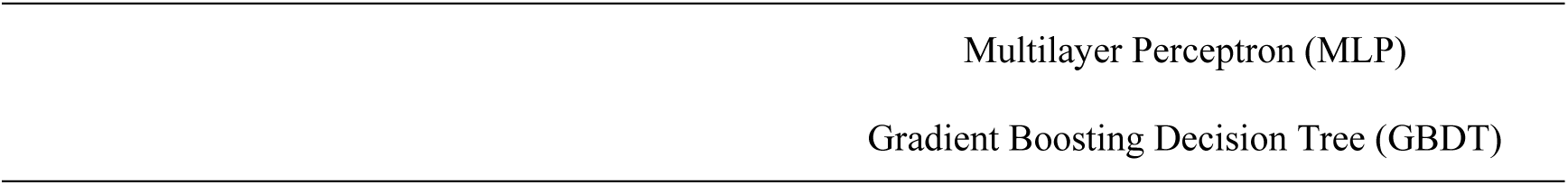
PLM embeddings and machine learning algorithms investigated during model.

**Table 3.** Model performance comparison of various PLM embeddings on training set and test set. The best results are highlighted in bold.

| Embedding | Model | Training set (CV=5) |  |  |  |  |  |  |  | Test set |  |  |  |  |  |  |  |
| --- | --- | --- | --- | --- | --- | --- | --- | --- | --- | --- | --- | --- | --- | --- | --- | --- | --- |
|  |  | Sn | Sp | PR | ACC | MCC | F1 | AU |  | Sn | Sp | PR | ACC | MCC | F1 | AU |  |
|  |  |  |  |  |  |  |  | ROC | PRC |  |  |  |  |  |  | ROC | PRC |
| ProtBert | MLP | **0.881** | 0.97 | 0.888 | 0.95 | **0.853** | **0.884** | **0.976** | 0.932 | **0.919** | 0.978 | 0.919 | **0.965** | **0.897** | **0.919** | **0.948** | 0.956 |
| ProtAlBert | MLP | 0.823 | 0.972 | 0.889 | 0.94 | 0.818 | 0.855 | 0.971 | 0.921 | 0.903 | 0.978 | 0.918 | 0.961 | 0.886 | 0.911 | 0.94 | **0.965** |
| ProtT5-XL-BFD | MLP | 0.819 | 0.973 | 0.892 | 0.94 | 0.817 | 0.854 | 0.953 | 0.917 | 0.855 | 0.978 | 0.914 | 0.951 | 0.853 | 0.883 | 0.916 | 0.949 |
| ProtBert-BFD | SVM | 0.798 | 0.973 | 0.89 | 0.935 | 0.803 | 0.842 | 0.956 | 0.9 | 0.855 | 0.973 | 0.898 | 0.947 | 0.843 | 0.876 | 0.914 | 0.897 |
| ESM-1v | Bagging | 0.774 | 0.974 | 0.891 | 0.931 | 0.788 | 0.828 | 0.945 | 0.899 | 0.839 | 0.987 | 0.945 | 0.954 | 0.863 | 0.889 | 0.913 | 0.904 |
| ProtT5-XL-UniRef50 | SVM | 0.823 | **0.986** | **0.943** | **0.951** | 0.852 | 0.879 | 0.97 | **0.943** | 0.823 | **1** | **1** | 0.961 | 0.885 | 0.903 | 0.911 | 0.943 |
| ESM-1b | LR | 0.823 | 0.97 | 0.881 | 0.938 | 0.813 | 0.851 | 0.959 | 0.93 | 0.839 | 0.982 | 0.929 | 0.951 | 0.852 | 0.881 | 0.91 | 0.959 |
| ESM-2-1280 | Adaboost | 0.778 | 0.98 | 0.913 | 0.936 | 0.805 | 0.84 | 0.943 | 0.903 | 0.823 | 0.987 | 0.944 | 0.951 | 0.852 | 0.879 | 0.905 | 0.951 |
| ESM-2-640 | SVM | 0.745 | 0.984 | 0.928 | 0.933 | 0.793 | 0.826 | 0.943 | 0.896 | 0.79 | 0.987 | 0.942 | 0.944 | 0.83 | 0.86 | 0.888 | 0.895 |
| ESM-2-480 | MLP | 0.753 | 0.945 | 0.789 | 0.904 | 0.71 | 0.771 | 0.944 | 0.879 | 0.79 | 0.991 | 0.961 | 0.947 | 0.841 | 0.867 | 0.891 | 0.925 |
| ESM-2-320 | LightGBM | 0.658 | 0.958 | 0.812 | 0.894 | 0.668 | 0.727 | 0.925 | 0.838 | 0.742 | 0.987 | 0.939 | 0.933 | 0.797 | 0.829 | 0.864 | 0.888 |

### 3.4 Comparative analysis of prediction models

The proposed models were further assessed and compared on the same test set. As observed, BERT-T6 show notable improvements across all metrics compared to the baseline model DDE_SVM, especially in Sn, MCC, F1-score and AUPRC, with an increase of approximately 32%, 29%, 24% and 19%, respectively (**Table 4**, **Figure 2**). Similarly, ProtBert_MLP also outperforms DDE_SVM on most metrics. That highlights the advantages of PLM, which excel at learning long-range residue interactions, capturing structural/functional semantics, and inferring evolutionary information. Compared to the ProtBert_MLP model, BERT-T6 achieves improvements in most of metrics, including ACC, Sn, F1-score, AUROC, MCC and AUPRC, with Sn, MCC and AUROC all showing the most significant increase (∼7%). This improvement is attributed to fine-tuning ProtBert on a T6SE-related dataset, which enhances its suitability for T6SE prediction. From another perspective, a key defect of predesigned feature representation methods is their inability to dynamically represent the semantic information of T6SEs[12].

**Table 4.** Comparative analysis of three representative classifiers proposed in this study across five independent test experiments. The best results are highlighted in bold.

| Model | ACC | Sn | Sp | PR | F1 | AUROC | MCC | AUPRC |
| --- | --- | --- | --- | --- | --- | --- | --- | --- |
| DDE_SV | 0.872 ± | 0.580 ± | 0.952 ± | 0.771 ± | 0.661 ± | 0.905 ± | 0.593 ± | 0.758 ± |
| M | 0.014 | 0.062 | 0.007 | 0.033 | 0.047 | 0.015 | 0.050 | 0.024 |
| ProtBert_ | 0.938 ± | 0.817 ± | 0.971 ± | 0.891 ± | 0.851 ± | 0.894 ± | 0.814 ± | 0.921 ± |
| MLP | 0.003 | 0.039 | 0.014 | 0.051 | 0.002 | 0.012 | 0.008 | 0.024 |
| BERT-T6 | **0.960 ±** | **0.902 ±** | **0.977 ±** | **0.916 ±** | **0.908 ±** | **0.973 ±** | **0.884 ±** | **0.950 ±** |
|  | **0.010** | **0.020** | **0.012** | **0.040** | **0.021** | **0.011** | **0.027** | **0.014** |

**Figure 2.**
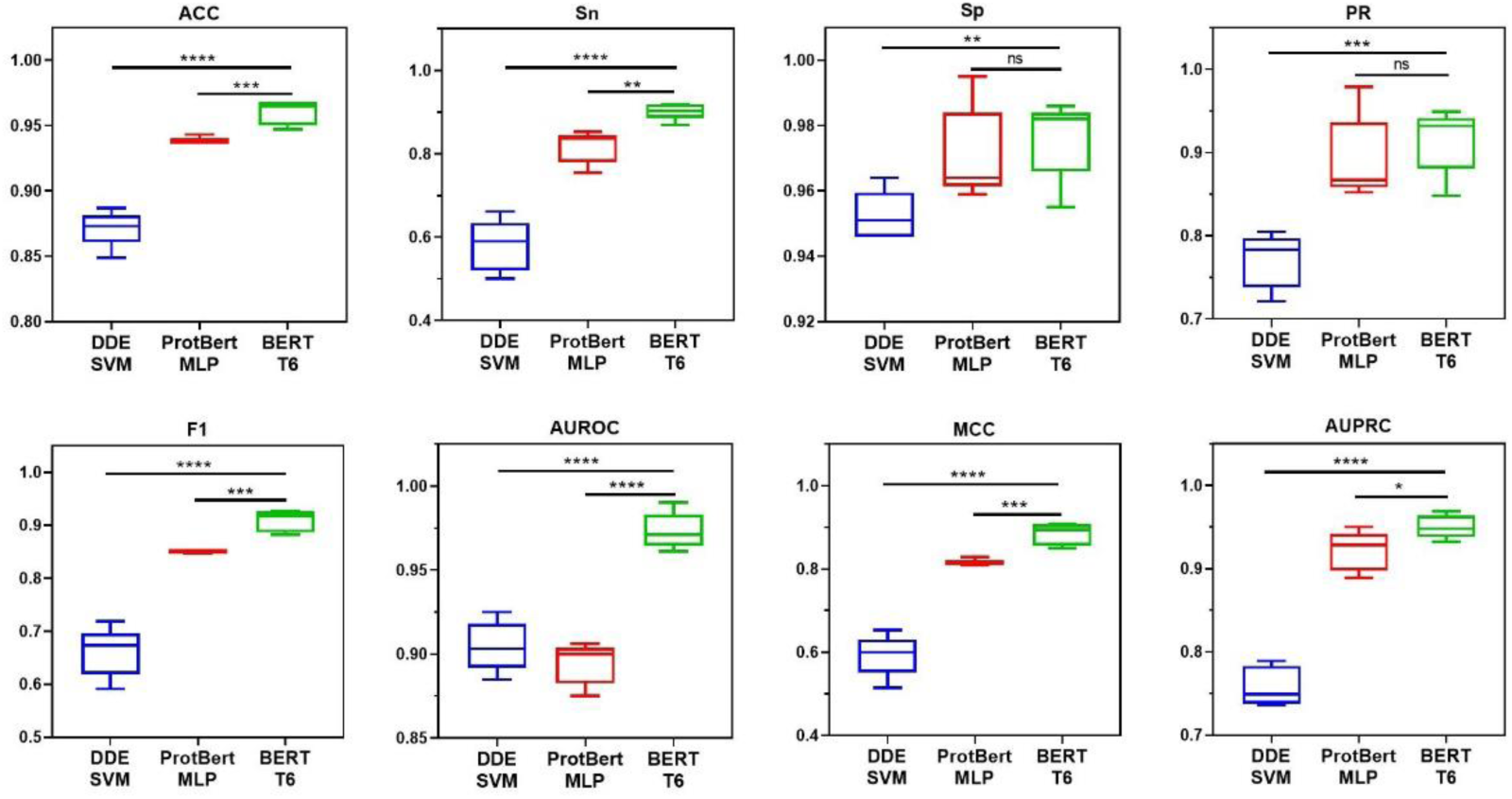
Performance comparison of proposed models on the test set across five repeated experiments.

### 3.5 Ablation study

To demonstrate the effectiveness of transfer learning in T6SE prediction, an ablation study was conducted. In this study, the ProtBert module was just used for feature extraction, and the fully-connected layers are kept the same as the architecture of BERT-T6, including loss function, optimizer, training epochs, dropout, learning rate and weight decay rate. The results are shown in **Table 5** and **Figure 3**, BERT-T6 successfully achieves significantly improved performance in five metrics based on the pretrained ProtBert model and the use of transfer learning techniques. This underlines, to some extent, the effectiveness of transfer learning for training robust predictive models in challenging classification tasks. On the contrary, without fine tune of ProtBert, there is a noticeable decline in the model’s performance, especially concerning is about the 22% decline in sensitivity and 14% decline in MCC. Additionally, F1-score, AUPRC and accuracy decreased by approximately 13%, 8.8% and 4%, respectively. That indicates features based on pre-trained ProtBert model cannot fully reveal specific sequential patterns of T6SEs though the model has been trained on a massive amount of protein sequence data. Employing a transfer learning strategy significantly enhanced the T6SE recognition performance of the model in terms of sensitivity. However, other evaluation metrics, such as specificity, precision, and AUROC, did not improve as significantly as expected, possibly due to the limited scale of the fine-tuning dataset. As more T6SEs are continuously discovered and the dataset expands, the impact of transfer learning on enhancing model performance will undoubtedly become more pronounced.

**Table 5.** Ablation Study. The best results are highlighted in bold.

| Model | Feature | ACC | Sn | Sp | PR | F1 | AUROC | MCC | AUPRC |
| --- | --- | --- | --- | --- | --- | --- | --- | --- | --- |
| MLP | ProtBert | 0.916 ± | 0.688 ± | 0.979 ± | 0.902 ± | 0.778 ± | 0.959 ± | 0.740 ± | 0.858 ± |
|  |  | 0.017 | 0.086 | 0.004 | 0.015 | 0.057 | 0.012 | 0.058 | 0.028 |
| BERT- | Embedding | **0.959 ±** | **0.909 ±** | **0.973 ±** | **0.905 ±** | **0.907 ±** | **0.970 ±** | **0.881 ±** | **0.946 ±** |
| T6 | layer | **0.011** | **0.009** | **0.015** | **0.046** | **0.023** | **0.014** | **0.031** | **0.019** |

**Figure 3.**
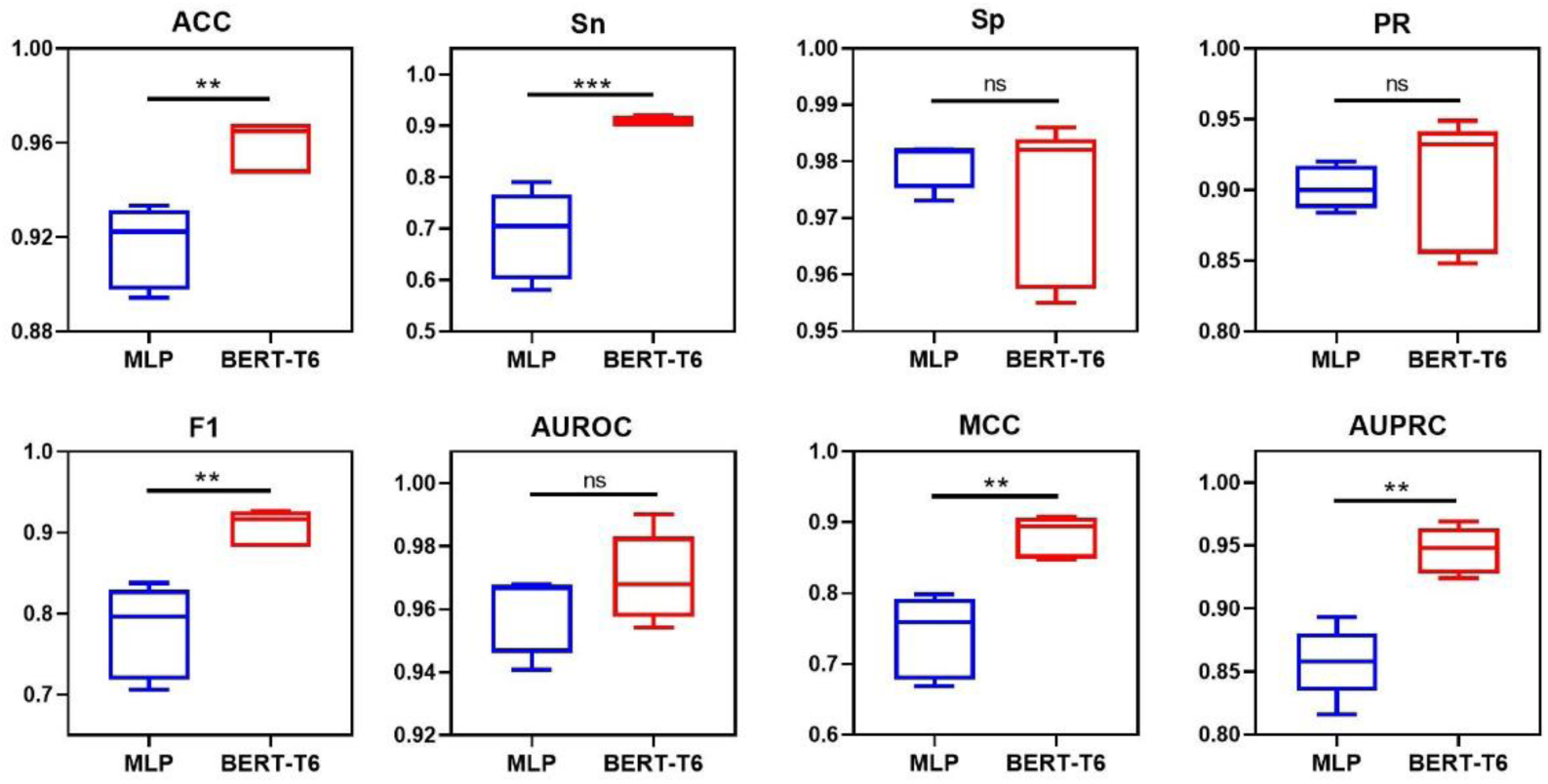
Ablation study.

**Figure 4.**
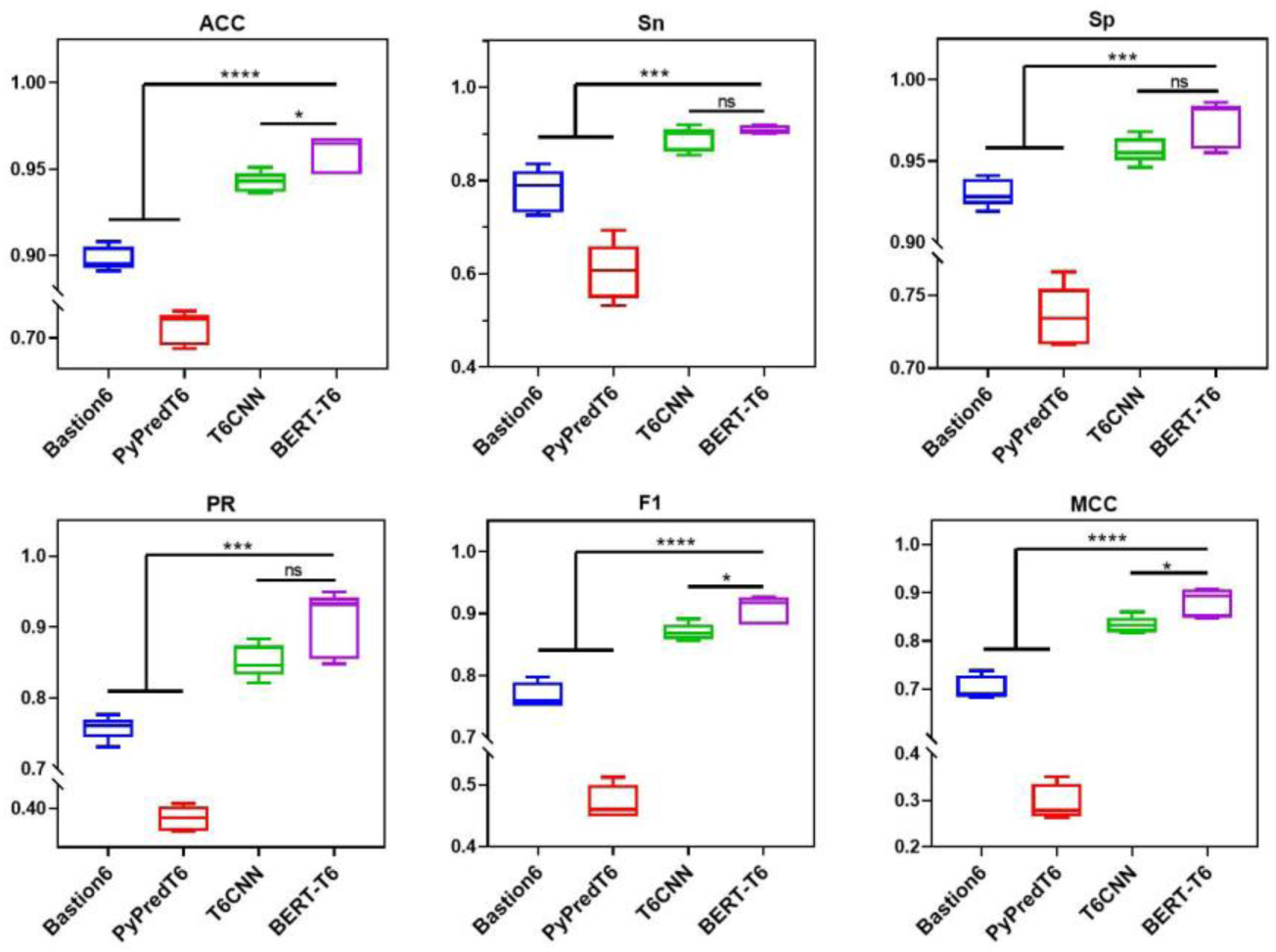
Performance of BERT-T6 and its comparison with SOTA models on the test set across five repeated experiments.

### 3.6 Performance comparison with existing tools

To validate the effectiveness of the proposed BERT-based transfer learning approach for T6SE prediction, we conducted the model performance comparison with existing tools on the test set across five repeated experiments. These existing tools includes Bastion6, PyPredT6 and T6CNN. Bastion6 is the first T6SE prediction tool, which is a two-layer SVM-based ensemble model integrating sequence-based, evolutionary information-based and physicochemical features[9]. Later, Sen et al.[42] designed PyPredT6 as a Python-based standalone tool which features are extracted from the peptide and nucleotide sequences of the experimentally verified T6SE, and it incorporates five algorithms including MLP, SVM, k-NN, Naive Bayes and RF, achieving prediction via majority voting. Recently, Hu et al.[10] proposed T6CNN, a CNN-based ensemble model for enhanced T6SE prediction, which integrates a diverse set of features, such as AAC-PSSM, EEDP, k-separated-bigrams-PSSM, Pse-PSSM, AADP-PSSM, an N-terminal 200-dimensional frequency matrix, and ESM-2. The experimental results are presented in **Table 5** and **Figure 4**. BERT-T6 outperforms Bastion6 and PyPredT6 across all six performance indicators. Specifically, our method achieves an accuracy of 0.959, a sensitivity of 0.909, a specificity of 0.973, a precision of 0.905, an F1-score of 0.907, and an MCC of 0.881. Compared to the second-best method, T6CNN, BERT-T6 improves accuracy, F1-score, and MCC by 1.7, 3.7, and 4.7%, respectively. While current SOTA models primarily rely on the integration of diverse manually selected features and an ensemble learning strategy, our approach achieves comparable or better results using only a single feature. This performance demonstrates the effectiveness of our fine-tuning strategy for the PLM ProtBert and underscores its potential advantages in T6SE prediction. The high sensitivity and specificity of BERT-T6 indicates its superior ability to distinguish T6SE and non T6SE compared to existing methods. Additionally, BERT-T6 achieves the highest MCC, demonstrating its overall classification effectiveness.

**Figure 4.**
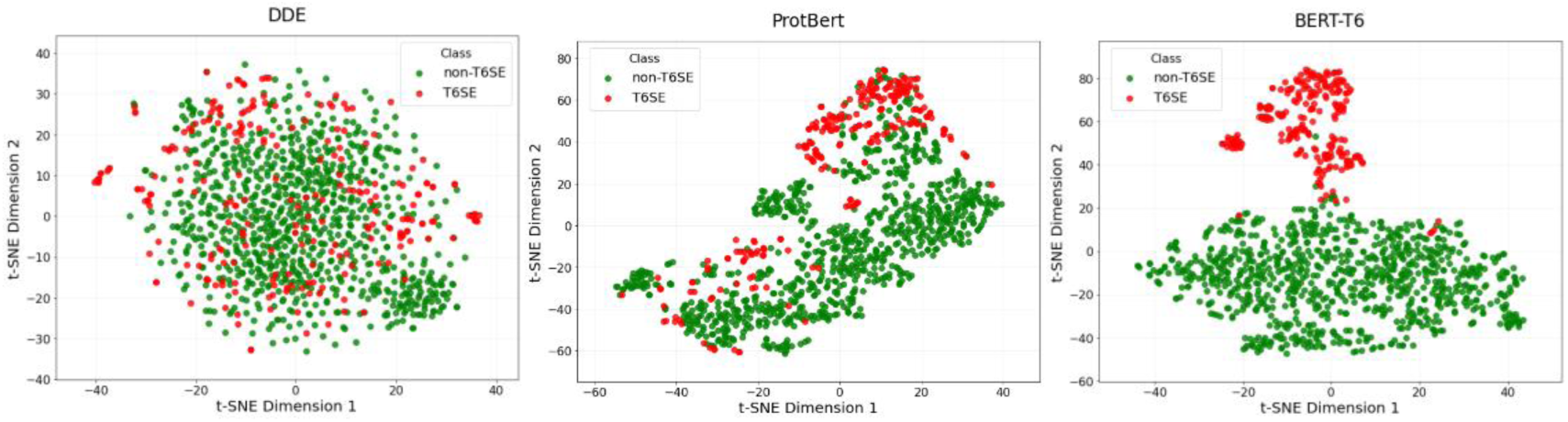
t-SNE visualization of the distribution of DDE, feature representations extracted from ProtBert and BERT-T6 embedding layer.

Besides the binary classifiers mentioned above, two multi-class models are involved in T6SE prediction: DeepSecE and CLEF. DeepSecE was proposed for the classification of T1SE, T2SE, T3SE, T4SE, T6SE, and non-secreted proteins. Its architecture contains a pre-trained ESM-2 module, a 1D convolutional layer, an additional transformer block, a fully connected layer, and a softmax activation function[43]. CLEF was developed for predicting T3SE, T4SE, and T6SE. It consists of two main modules: a contrastive learning module that encodes protein sequence representations into cross-modality representations aligned with biological features, and an effector prediction module that utilizes these representations to predict effectors[44]. Here, we present a comparative evaluation of BERT-T6 against these two models. As presented in **Table S5** and **Figure S1**, BERT-T6 achieves a higher sensitivity than DeepSecE and CLEF. Although its precision is relatively lower than that of DeepSecE and CLEF, BERT-T6 exhibits comparable performance in accuracy, F1-score, and MCC. In conclusion, the proposed model exhibits a significant advantage over existing binary classifiers and competitive performance compared to SOTA multi-class models, making it a promising approach for T6SE prediction.

**Table 5.**
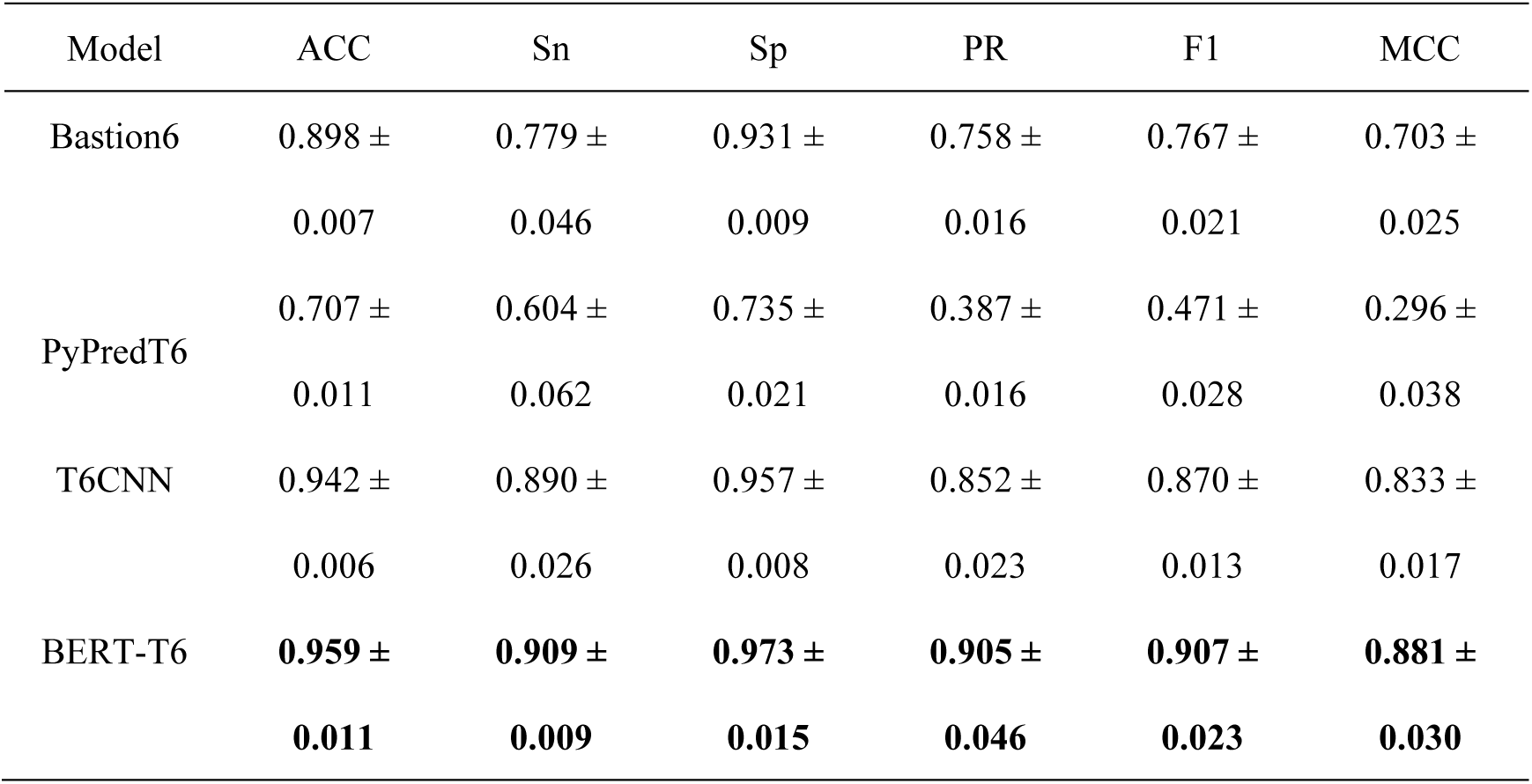
Performance comparison of BERT-T6 with existing SOTA models on the test set. The best results are highlighted in bold.

| Model | ACC | Sn | Sp | PR | F1 | MCC |
| --- | --- | --- | --- | --- | --- | --- |
| Bastion6 | 0.898 ± 0.007 | 0.779 ± 0.046 | 0.931 ± 0.009 | 0.758 ± 0.016 | 0.767 ± 0.021 | 0.703 ± 0.025 |
| PyPredT6 | 0.707 ± 0.011 | 0.604 ± 0.062 | 0.735 ± 0.021 | 0.387 ± 0.016 | 0.471 ± 0.028 | 0.296 ± 0.038 |
| T6CNN | 0.942 ± 0.006 | 0.890 ± 0.026 | 0.957 ± 0.008 | 0.852 ± 0.023 | 0.870 ± 0.013 | 0.833 ± 0.017 |
| BERT-T6 | **0.959 ± 0.011** | **0.909 ± 0.009** | **0.973 ± 0.015** | **0.905 ± 0.046** | **0.907 ± 0.023** | **0.881 ± 0.030** |

### 3.7 Performance validation on the independent test set

To validate the model’s generalization ability, we further evaluated BERT-T6 and performed comparison with SOTA models on the independent test. As exhibited in **Table 6**, the BERT-T6 achieves the highest score for accuracy (ACC = 0.896), sensitivity (Sn = 0.966), F1-score (0.809) and MCC (0.758), while T6CNN achieves the highest score for accuracy (ACC = 0.896), specificity (Sp = 0.900) and precision (PR = 0.722). Compared to the T6CNN model, although a slight decrease in specificity (Sp = 0.875) and precision (PR = 0.695) is observed, the BERT-T6 model achieves a substantially higher sensitivity. This superior performance highlights its capability to identify T6SEs from unknown samples. Furthermore, the model achieves a relative balance between sensitivity and specificity from the prediction of unbalanced samples. In addition, we compared BERT-T6 with two multi-classifiers on an independent test set across the five metrics. The results are presented in **Table S6**. Our model achieves the highest scores in accuracy and sensitivity, and the second highest in F1-score and MCC among the three models. In summary, these results collectively demonstrate the usefulness and strong generalization capability of BERT-based transfer learning approach for T6SE prediction on unseen data.

**Table 6.** Performance comparison of BERT-T6 with existing SOTA models on independent test set. The best results are highlighted in bold.

| Model | ACC | Sn | Sp | PR | F1 | MCC |
| --- | --- | --- | --- | --- | --- | --- |
| Bastion6 | 0.846 | 0.712 | 0.885 | 0.646 | 0.677 | 0.577 |
| PyPredT6 | 0.730 | 0.542 | 0.785 | 0.427 | 0.478 | 0.303 |
| T6CNN | **0.896** | 0.881 | **0.900** | **0.722** | 0.794 | 0.731 |
| BERT-T6 | **0.896** | **0.966** | 0.875 | 0.695 | **0.809** | **0.758** |

### 3.8 Interpretable analysis

#### 3.8.1 Visualizing the features learned by BERT-T6 after fine tune of ProtBert

To intuitively estimate the effect of different feature encoding methods on the classification performance as well as acquire an intuitive understanding of the features learned by BERT-T6 after fine tune of ProtBert model, we applied the t-distributed stochastic neighbor embedding (t-SNE) to map all the samples onto two-dimensional space. As shown in **Figure 4**, DDE-based sequence features fail to adequately discriminate T6SE from non-T6SE due to overlapping clusters formed by these features. In contrast, ProtBert embedding-based features demonstrate improved performance, with clusters of positive and negative samples achieving near-separation, though partial overlap is observed in the lower-left and upper-right corners. Notably, BERT-T6 can learn even more effective feature representations than pretrained ProtBert embeddings, as evidenced by the more distinct clustering of samples from the same category and better separation between samples of different categories. It demonstrates that the model, through training, successfully extracts and differentiates the key features of samples from different classes, highlighting its exceptional classification capability.

#### 3.8.2 Case study with attention analysis

In addition to the t-SNE analysis, we conducted a detailed examination of the model’s attention mechanisms. Attention mechanisms provide insights into which parts of the input data the model focuses on during the classification process. By analyzing these attention maps, we could identify how the model emphasizes different regions of the data and how these regions impact the classification results, which help us get a better understanding and put more trust on this model. The first case is the Ntox15 family nuclease effector (Tde1), which mediates interbacterial competition among Bacteroidales by degrading recipient genomic DNA via its Ntox15 DNase domain[23]. The sequence of Tde1 was input into the BERT-T6 model, and the attention score matrices from the 16 heads of the last layer in the ProtBert module were extracted and summed. The amino acid-level attention scores were then calculated by summing the columns, which was visualized as **Figure 5**. Here, the value of attention scores greater than the interquartile range are regarded as significant attention scores and highlighted with gray box[45]. As observed, approximately 92% of amino acids with significant attention scores are located within the Ntox15 domain, which means that influence the most in the model prediction, even though this domain covers only 43% of the total sequence length of Tde1. This reveals BERT-T6 model pay more attention on the function region of Tde1 after training.

**Figure 5.**
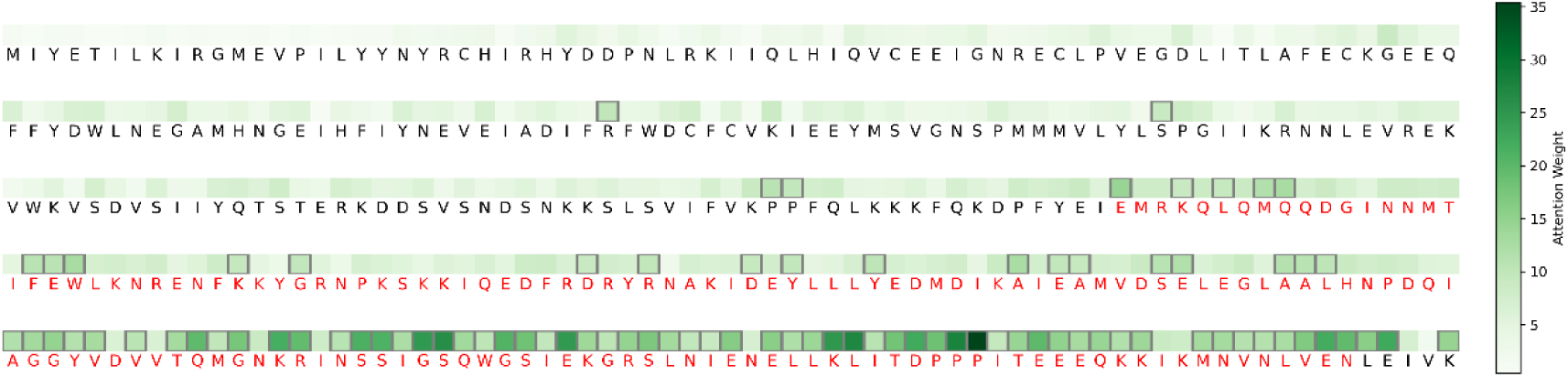
Attention heatmap for Tde1. The amino acids sequence labeled in red color represents Ntox15 domain of Tde1. The gray box highlights the more significant attention scores.

The second case is AvPCP, an effector protein produced by *Anabaena variabilis*. Unlike the cytoplasmic effector Tde1, AvPCP is injected into the periplasm of recipient cells. Its NlpC/P60 domain acts as a γ-D-glutamyl-L-diamino acid endopeptidase, catalyzing the hydrolysis of the peptide bond between γ-D-glutamyl and L-diamino acid residues in peptidoglycan[46]. A similar analysis was conducted on the sequence of AvPCP, and the results are presented in **Figure 6**. Approximately 81% of amino acids with significant attention scores are located within the NlpC/P60 domain, while the NlpC/P60 domain only occupies 47% of the total length. This indicates that these residues play an important role in T6SEs prediction. In total, these analyses demonstrate that through fine-tuning, BERT-T6 learned to identify functional regions in given T6SEs that contribute to its overall binary classification performance.

**Figure 6.**
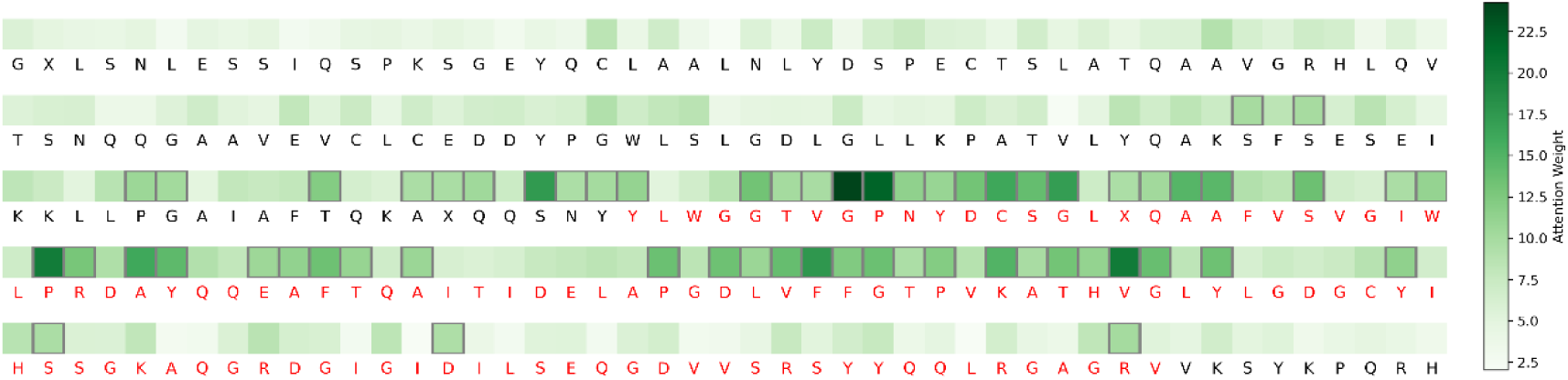
Attention heatmap for AvPCP. The amino acids sequence labeled in red color represents NlpC/P60 domain of AvPCP. The gray box highlights the more significant attention scores.

## 4. Conclusion

In this study, we successively investigated two types of features, namely those based on sequence-encoding and those based on PLM embedding, for use in machine learning modeling of T6SE prediction. DDE represents the optimal sequence-based feature to build the model when integrates with the SVM algorithm. However, lower sensitivity (∼60%) and F1-score make it difficult to recognize T6SE accurately. In contrast, the use of features based on PLM embeddings significantly improves the sensitivity and robustness of T6SE prediction. In particular, the model with embeddings derived from the ProtBert pre-trained PLM as input performs the best on the test set, achieving the highest sensitivity, accuracy, MCC, F1-score, and AUROC. Based on that, we further fine-tune ProtBert model through downstream T6SE classification task, and the resulting BERT-T6 model has improved performance in most evaluation metrics. Encouragingly, our model exhibits the superior performance compared to the previous SOTA tools in terms of accuracy, sensitivity, F1-score and MCC. In total, BERT-based transfer learning approach is demonstrated as an effective strategy for the improvement of T6SE prediction. Combined with interpretability analysis via the attention mechanism, our model not only predicts T6SEs efficiently but also accurately identifies and focuses on key functional regions within the sequences. This provides biologists with a powerful tool to gain deeper insights into the relationship between protein sequences and toxicity. Overall, BERT-T6 serves as a precise, robust, and interpretable solution that accelerates the identification and characterization of novel T6SEs. Future work will focus on exploitation of dual-task protein language pretrained model for simultaneous T6SE identification and categorization prediction.

## Acknowledgments

This project was funded by Macao Polytechnic University with grant number RP/FCA-06/2024. The funders had no role in study design, data collection, and interpretation, or the decision to submit the work for publication. J. C. is recipient of the Macau Polytechnic University (MPU) graduate scholarship. This project is part of the thesis work of X. M. with an internal reference number of s/c xxxx.

## Data Availability

All T6SEs and non-T6SEs data used in this study comes from Bastion6 (https://bastion6.erc.monash.edu/download.jsp), SecReT6 (https://bioinfomml.sjtu.edu.cn/SecReT6/index.php) and TxSEdb (http://61.160.194.165:3080/TxSEdb/index.html). The code used to develop the model, perform T6SE prediction are publicly available in GitHub (https://github.com/mxw1992/BERT-T6).

