## Supplementary Material for "BERT-T6: Towards High-accuracy T6SS Bacterial Toxin Identification Using Protein Language Model"

**Table S1 Comparison of model performance with various sequence encoding methods on training set, sorted by AUROC.** The best results are highlighted in bold, while the second-best results are underlined.

| Model | Training set (5CV) | | | | | | | |
|---|---|---|---|---|---|---|---|---|
| | Sensitivity (%) | Specificity (%) | Precision (%) | Accuracy (%) | MCC | F1 | AUROC | AUPRC |
| DDE_SVM_model | **51.42** | 93.574 | 69.988 | **84.512** | **0.5075** | **0.5874** | **0.8771** | **0.7002** |
| QSOrder_SVM_model | 42.74 | 93.574 | 65.418 | 82.658 | 0.4295 | 0.5129 | 0.8576 | 0.6146 |
| CTDC_SVM_model | 29.176 | 94.926 | 62.442 | 80.796 | 0.3274 | 0.3881 | 0.816 | 0.5433 |
| KSCTriad_SVM_model | 27.178 | 95.606 | 62.566 | 80.884 | 0.3198 | 0.3753 | 0.775 | 0.5322 |
| CTriad_SVM_model | 23.86 | 96.394 | 64.818 | 80.796 | 0.3071 | 0.3453 | 0.7641 | 0.5109 |
| CTDT_SVM_model | 0 | **100** | 0 | 78.496 | 0 | 0 | 0.7638 | 0.448 |
| PAAC_SVM_model | 9.446 | 99.888 | **97.5** | 80.442 | 0.2658 | 0.1706 | 0.7565 | 0.5216 |
| APAAC_SVM_model | 2.048 | **100** | 60 | 78.936 | 0.0975 | 0.0395 | 0.7492 | 0.5156 |
| NMBroto_SVM_model | 0 | **100** | 0 | 78.496 | 0 | 0 | 0.7391 | 0.4429 |
| GAAC_SVM_model | 0 | 98.99 | 0 | 77.7 | -0.0325 | 0 | 0.7265 | 0.4013 |
| CTDD_SVM_model | 3.69 | 99.212 | 44 | 78.672 | 0.0878 | 0.0677 | 0.5786 | 0.3326 |
| SOCNumber_SVM_model | 0 | **100** | 0 | 78.496 | 0 | 0 | 0.5 | 0.6075 |
| Moran_SVM_model | 0 | **100** | 0 | 78.496 | 0 | 0 | 0.3609 | 0.1628 |
| Geary_SVM_model | 0 | **100** | 0 | 78.496 | 0 | 0 | 0.3589 | 0.1624 |
| GTPC_SVM_model | 0 | **100** | 0 | 78.496 | 0 | 0 | 0.2924 | 0.1471 |
| CKSAAGP_SVM_model | 0 | **100** | 0 | 78.496 | 0 | 0 | 0.2799 | 0.1447 |
| GDPC_SVM_model | 0 | **100** | 0 | 78.496 | 0 | 0 | 0.2749 | 0.1437 |
| AAC_SVM_model | 0 | **100** | 0 | 78.496 | 0 | 0 | 0.1813 | 0.1297 |
| DistancePair_SVM_model | 0 | **100** | 0 | 78.496 | 0 | 0 | 0.1813 | 0.1297 |
| ASDC_SVM_model | 0 | **100** | 0 | 78.496 | 0 | 0 | 0.1774 | 0.1291 |

| | | | | | | | | |
|---|---|---|---|---|---|---|---|---|
| TPC_SVM_model | 0 | **100** | 0 | 78.496 | 0 | 0 | 0.1759 | 0.1287 |
| DPC_SVM_model | 0 | **100** | 0 | 78.496 | 0 | 0 | 0.1731 | 0.1287 |
| CKSAAP_SVM_model | 0 | **100** | 0 | 78.496 | 0 | 0 | 0.1685 | 0.1282 |

**Table S2 Comparison of model performance with various sequence encoding methods on test set, sorted by AUROC.** The best results are highlighted in bold, while the second-best results are underlined.

| Model | Test_set | | | | | | | |
|---|---|---|---|---|---|---|---|---|
| | Sensitivity (%) | Specificity (%) | Precision (%) | Accuracy (%) | MCC | F1 | AUROC | AUPRC |
| DDE_SVM_model | **66.13** | 95.07 | 78.85 | **88.77** | **0.6537** | **0.7193** | **0.8992** | **0.7882** |
| QSOrder_SVM_model | 48.39 | 95.07 | 73.17 | 84.91 | 0.5109 | 0.5825 | 0.8555 | 0.6561 |
| CTDC_SVM_model | 40.32 | 95.07 | 69.44 | 83.16 | 0.4395 | 0.5102 | 0.8187 | 0.5951 |
| PAAC_SVM_model | 1.61 | **100** | **100** | 78.6 | 0.1125 | 0.0317 | 0.8133 | 0.5165 |
| CTriad_SVM_model | 19.35 | 99.1 | 85.71 | 81.75 | 0.3524 | 0.3158 | 0.7997 | 0.6009 |
| APAAC_SVM_model | 1.61 | **100** | 100 | 78.6 | 0.1125 | 0.0317 | 0.7992 | 0.5103 |
| KSCTriad_SVM_model | 25.81 | 97.76 | 76.19 | 82.11 | 0.3721 | 0.3855 | 0.7908 | 0.5879 |
| CTDT_SVM_model | 0 | **100** | 0 | 78.25 | 0 | 0 | 0.7854 | 0.4443 |
| NMBroto_SVM_model | 0 | **100** | 0 | 78.25 | 0 | 0 | 0.7642 | 0.5154 |
| GAAC_SVM_model | 0 | 99.55 | 0 | 77.89 | -0.0313 | 0 | 0.6881 | 0.3713 |
| CTDD_SVM_model | 14.52 | **100** | **100** | 81.4 | 0.3425 | 0.2535 | 0.6667 | 0.5159 |
| SOCNumber_SVM_model | 0 | **100** | 0 | 78.25 | 0 | 0 | 0.4919 | 0.6011 |
| Geary_SVM_model | 0 | **100** | 0 | 78.25 | 0 | 0 | 0.3263 | 0.1538 |
| GTPC_SVM_model | 0 | **100** | 0 | 78.25 | 0 | 0 | 0.3261 | 0.1565 |
| Moran_SVM_model | 0 | **100** | 0 | 78.25 | 0 | 0 | 0.3185 | 0.1519 |
| CKSAAGP_SVM_model | 0 | **100** | 0 | 78.25 | 0 | 0 | 0.3104 | 0.1541 |
| GDPC_SVM_model | 0 | **100** | 0 | 78.25 | 0 | 0 | 0.2998 | 0.1521 |
| TPC_SVM_model | 0 | **100** | 0 | 78.25 | 0 | 0 | 0.1854 | 0.1304 |
| DPC_SVM_model | 0 | **100** | 0 | 78.25 | 0 | 0 | 0.1784 | 0.1302 |
| AAC_SVM_model | 0 | **100** | 0 | 78.25 | 0 | 0 | 0.1783 | 0.1304 |
| DistancePair_SVM_model | 0 | **100** | 0 | 78.25 | 0 | 0 | 0.1783 | 0.1304 |
| ASDC_SVM_model | 0 | **100** | 0 | 78.25 | 0 | 0 | 0.1735 | 0.1301 |
| CKSAAP_SVM_model | 0 | **100** | 0 | 78.25 | 0 | 0 | 0.1704 | 0.1293 |

**Table S3 Comparison of model performance with various machine learning algorithms on training set, sorted by AUROC.** The best results are highlighted in bold, while the second-best results are underlined.

| Model (DDE) | Training set (5CV) | | | | | | | |
|---|---|---|---|---|---|---|---|---|
| | Sensitivity (%) | Specificity (%) | Precision (%) | Accuracy (%) | MCC | F1 | AUROC | AUPRC |
| **SVM_model** | 51.42 | 93.574 | 69.988 | **84.512** | **0.5075** | 0.5874 | **0.8771** | **0.7002** |
| LightGBM_model | 34.566 | 97.52 | **80.14** | 83.982 | 0.4543 | 0.481 | 0.8462 | 0.6446 |
| NaiveBayes_model | **64.582** | 85.34 | 54.628 | 80.884 | 0.471 | **0.5914** | 0.8364 | 0.5988 |
| MLP_model | 53.48 | 91.882 | 64.09 | 83.628 | 0.4846 | 0.5787 | 0.8345 | 0.6223 |
| XGBoost_model | 36.616 | 96.728 | 75.894 | 83.806 | 0.4483 | 0.4899 | 0.8329 | 0.6449 |
| GBDT_model | 29.616 | 96.054 | 67.412 | 81.77 | 0.3602 | 0.4103 | 0.8168 | 0.5834 |
| Adaboost_model | 47.68 | 92.22 | 62.58 | 82.654 | 0.442 | 0.5375 | 0.8143 | 0.5796 |
| RF_model | 25.908 | **98.196** | 79.444 | 82.656 | 0.3865 | 0.389 | 0.8058 | 0.608 |
| LR_model | 51.004 | 85.57 | 49.038 | 78.144 | 0.3602 | 0.4989 | 0.7764 | 0.5053 |
| SGD_model | 51.014 | 84.556 | 47.526 | 77.346 | 0.3468 | 0.49 | 0.7611 | 0.5398 |
| KNN_model | 37.406 | 96.052 | 73.726 | 83.45 | 0.4407 | 0.4906 | 0.7506 | 0.6145 |
| Bagging_model | 34.134 | 92.784 | 56.74 | 80.178 | 0.3296 | 0.4247 | 0.7349 | 0.4682 |
| LDA_model | 51.386 | 82.414 | 44.314 | 75.754 | 0.3207 | 0.4755 | 0.7318 | 0.4552 |
| DT_model | 40.316 | 82.858 | 39.398 | 73.716 | 0.2303 | 0.3969 | 0.6159 | 0.4627 |

**Table S4 Comparison of model performance with various machine learning algorithms on test set, sorted by AUROC.** The best results are highlighted in bold, while the second-best results are underlined**.**

| Model (DDE) | Test_set | | | | | | | |
|---|---|---|---|---|---|---|---|---|
| | Sensitivity (%) | Specificity (%) | Precision (%) | Accuracy (%) | MCC | F1 | AUROC | AUPRC |
| SVM_model | 66.13 | 95.05 | 78.85 | **88.73** | **0.6534** | **0.7193** | **0.8992** | **0.7885** |
| NaiveBayes_model | **70.97** | 90.54 | 67.69 | 86.27 | 0.6048 | 0.6929 | 0.88 | 0.7031 |
| LightGBM_model | 38.71 | 98.65 | 88.89 | 85.56 | 0.5262 | 0.5393 | 0.8633 | 0.7321 |
| XGBoost_model | 40.32 | 97.3 | 80.65 | 84.86 | 0.4984 | 0.5376 | 0.8632 | 0.7076 |
| MLP_model | 54.84 | 93.24 | 69.39 | 84.86 | 0.5257 | 0.6126 | 0.8602 | 0.6867 |
| RF_model | 32.26 | 99.1 | 90.91 | 84.51 | 0.4846 | 0.4762 | 0.8595 | 0.7368 |
| Bagging_model | 27.42 | **99.55** | **94.44** | 83.8 | 0.4573 | 0.425 | 0.8588 | 0.6939 |
| GBDT_model | 33.87 | 98.65 | 87.5 | 84.51 | 0.483 | 0.4884 | 0.858 | 0.7158 |
| SGD_model | 61.29 | 87.39 | 57.58 | 81.69 | 0.4761 | 0.5938 | 0.8372 | 0.6141 |
| LR_model | 56.45 | 88.74 | 58.33 | 81.69 | 0.4573 | 0.5738 | 0.8356 | 0.6225 |
| Adaboost_model | 48.39 | 92.79 | 65.22 | 83.1 | 0.4617 | 0.5556 | 0.8325 | 0.6498 |
| KNN_model | 41.94 | 97.3 | 81.25 | 85.21 | 0.5126 | 0.5532 | 0.8247 | 0.6857 |
| DT_model | 41.94 | 93.69 | 65 | 82.39 | 0.4231 | 0.5098 | 0.7992 | 0.6108 |
| LDA_model | 53.23 | 86.04 | 51.56 | 78.87 | 0.3882 | 0.5238 | 0.7864 | 0.6053 |

**Table S5 Performance comparison of BERT-T6 with existing SOTA multi-classifiers on test set.** The best results are highlighted in bold.

| Model | ACC | Sn | PR | F1 | MCC |
|---|---|---|---|---|---|
| DeepSecE | 0.950 ± 0.007 | 0.818 ± 0.046 | 0.984 ± 0.016 | 0.893 ± 0.031 | 0.873 ± 0.033 |
| CLEF | **0.960 ± 0.009** | 0.857 ± 0.038 | **1.000 ± 0.000** | **0.923 ± 0.022** | **0.908 ± 0.025** |
| BERT-T6 | 0.959 ± 0.011 | **0.909 ± 0.009** | 0.905 ± 0.046 | 0.907 ± 0.023 | 0.881 ± 0.030 |

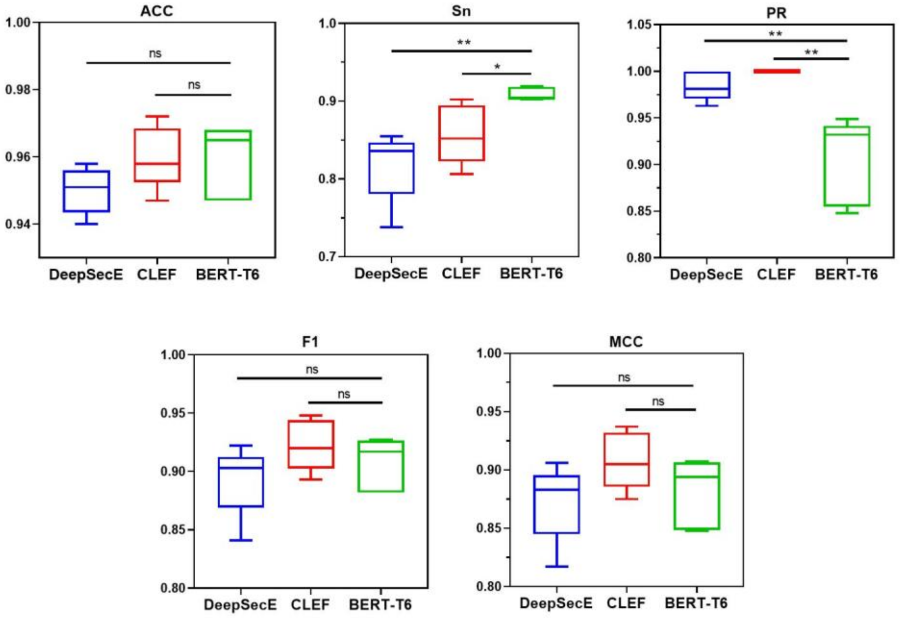


**Figure S1 Performance of BERT-T6 and its comparison with SOTA multi-classifiers on the test set across five repeated experiments.**

**Table S6 Performance comparison of BERT-T6 with existing SOTA multi-classifiers on independent test set.** The best results are highlighted in bold, while the second-best results are underlined.

| Model | ACC | Sn | PR | F1 | MCC |
|---|---|---|---|---|---|
| DeepSecE | 0.869 | 0.746 | **1.000** | **0.855** | **0.830** |
| CLEF | 0.849 | 0.661 | 0.975 | 0.788 | 0.758 |
| BERT-T6 | **0.896** | **0.966** | 0.695 | 0.809 | 0.758 |
